# Analysis and Engineering of Substrate Shuttling by the Acyl Carrier Protein (ACP) in Fatty Acid Synthases (FASs)

**DOI:** 10.1101/371880

**Authors:** Emanuele Rossini, Jan Gajewski, Maja Klaus, Gerhard Hummer, Martin Grininger

## Abstract

In the large enzyme complexes of natural biosynthetic pathways, molecules are assembled like in a factory. Carrier domains shuttle substrates and intermediates as covalently attached cargo within the enzyme complex between active sites. The physical confinement of the reaction increases reaction rates and hinders pathway branching. Alternating interactions of substrate-loaded carrier domains with different catalytic domains modulate the chemical environment. In this study, we aim at assessing the impact of domain-domain interactions (DDIs) on the reaction progress of a multienzyme type I fatty acid synthase (FAS) in quantitative terms. We modulate DDIs by single interface mutations, and read out the impact on substrate shuttling by recording fatty acid (FA) chain length product spectra and FAS activities. Our data show that even single interface point mutations can severely affect FA synthesis. With molecular dynamics simulations and modeling, we relate the mutation effects to specific alterations in the molecular interaction networks and domain-domain binding energetics. Some of the presented mutations induce the synthesis of short-chain FAs. These compounds are important commodity products and potent precursors for microbial biofuel production.

---

Living cells produce fatty acids (FA) in an iterative yet tightly controlled process (Fig. 1A).^1–4^ As the decisive step in chain length control by fatty acid synthase (FAS), the fully reduced acyl chain, bound to the acyl carrier protein (ACP) (Fig. 1B),^5^ is either transferred to the condensing ketoacyl synthase (KS) for another elongation, or released by a transferase (MPT) or a thio-esterase (TE) (Fig. SI1).^6–8^ In bacterial and fungal multi-enzyme (type I) FASs, chain length is controlled by the competitive substrate acceptance of the KS domain and the MPT domain, and as such is influenced by the interfaces ACP:KS and ACP:MPT.^9–10^ We recently engineered the G2559S-M2600W-mutated *Corynebacterium ammoniagenes* FAS, hereafter termed FAS^GSMW^, to produce a bimodal spectrum of C_8_- and C_14_/C_16_-CoA *in vitro*.^11–12^ The two mutations of FAS^GSMW^ are located in the KS-binding channel, where they attenuate the loading of octanoyl in the KS binding channel and promote the off-loading of the acyl chain by the substrate-promiscuous MPT to produce C_8_-CoA. With its bimodal product spectrum, the FAS^GSMW^ mutant is a well-suited reporter system for elaborating the impact of domain-domain interactions (DDIs) in substrate shuttling (Fig. 1C). We expected shifts in the proportion of C_8_- and C_14_/C_16_-CoA by perturbing the interfaces between the carrier protein ACP with domains KS (ACP:KS) and MPT (ACP:MPT). Mutations decreasing the affinity between the ACP and KS domains should steer synthesis towards short-chain products by promoting the release from the MPT domain. Conversely, increased ACP:KS affinity should decrease the share of short-chain products. In the following, we report data collected on the ACP:KS interface. For ACP:MPT interface engineering, see Note SI1 and SI Fig. SI2 and SI3.

**Figure 1.**
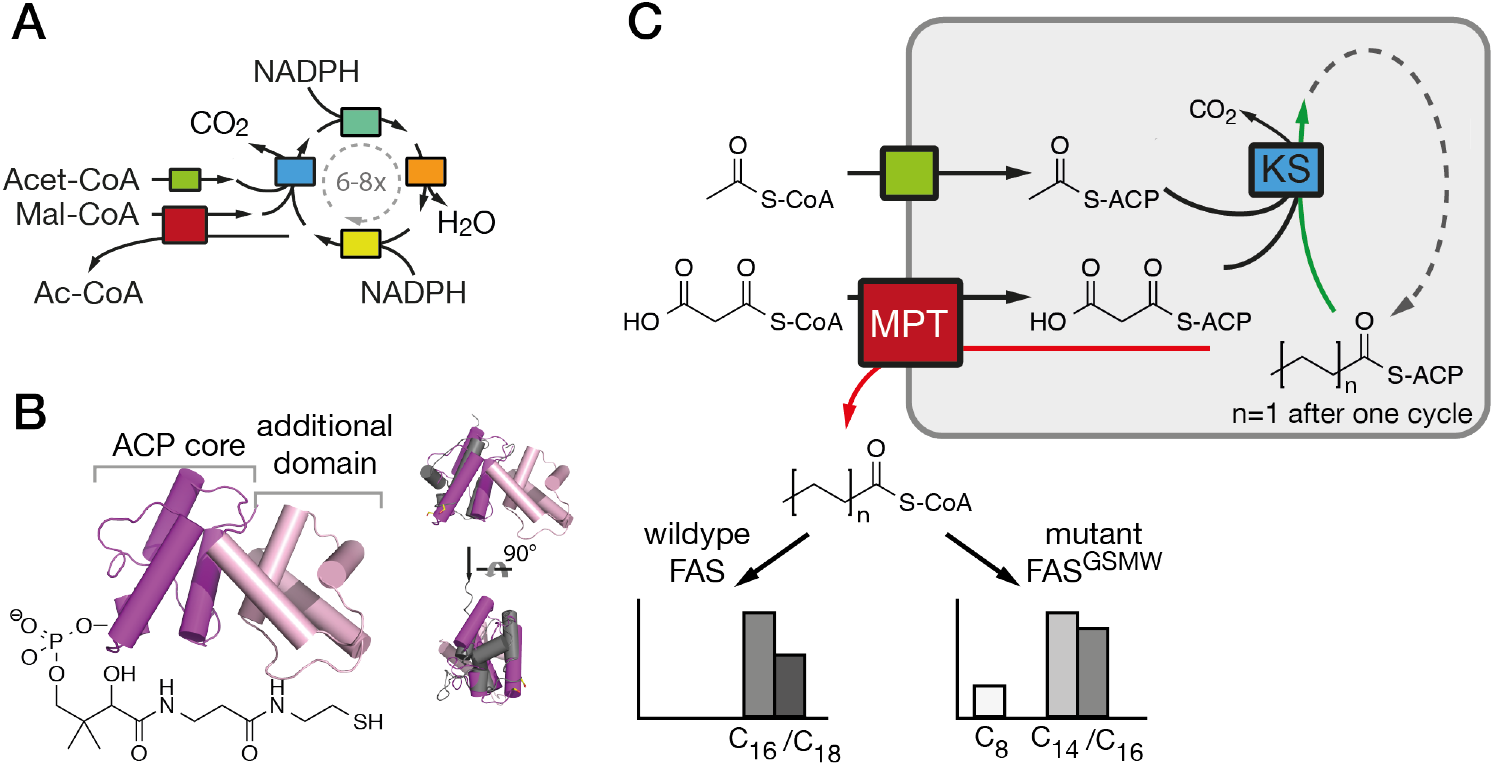
FA synthesis. A) Iterative type I bacterial and fungal FA synthesis produces acyl-CoA esters. (B) Phosphopantetheinylated ACPs covalently binds substrates and intermediates. ACPs of type I bacterial and fungal FASs are comprised of two domains (yeast FAS ACP magenta; pdb 2uv8^17^), in contrast to other ACPs (rat FAS ACP in grey; 2png^25^). (C) Modulated chain length regulation in FAS^GSMW^ induces a bimodal product spectrum.

In light of the need for microbial short-chain FA production,^12–14^ we aimed at mutating the ACP:KS interface to shift the FAS^GSMW^ product spectrum towards short-chain products. To modulate the ACP:KS interface, mutations were introduced on the surface of the KS domain only, since an intact ACP leaves other DDIs during synthesis unaffected (Fig. 2A). Mutation sites on the KS domain surface, such as D2553, D2556, and N2557, were derived from a study analyzing the transient interaction of FabB, a KS homologous protein in *E. coli*,^15^ and its ACP. A2696 was identified from a crosslinking study on FabF, another KS homologue in *E. coli*.^16^ Additionally, structural data of the *S. cerevisiae* FAS, which was crystallized with ACP docked to the KS domain,^17–18^ were included in our consideration. On this basis, we selected two novel interaction sites, N2621 and D2622 (Fig. 2A and Table SI1). The selected residues were mutated to alanine and/or to residues with opposite charges, and the mutant protein analyzed in activity and product spectra (SI Material and Methods).

**Figure 2.**
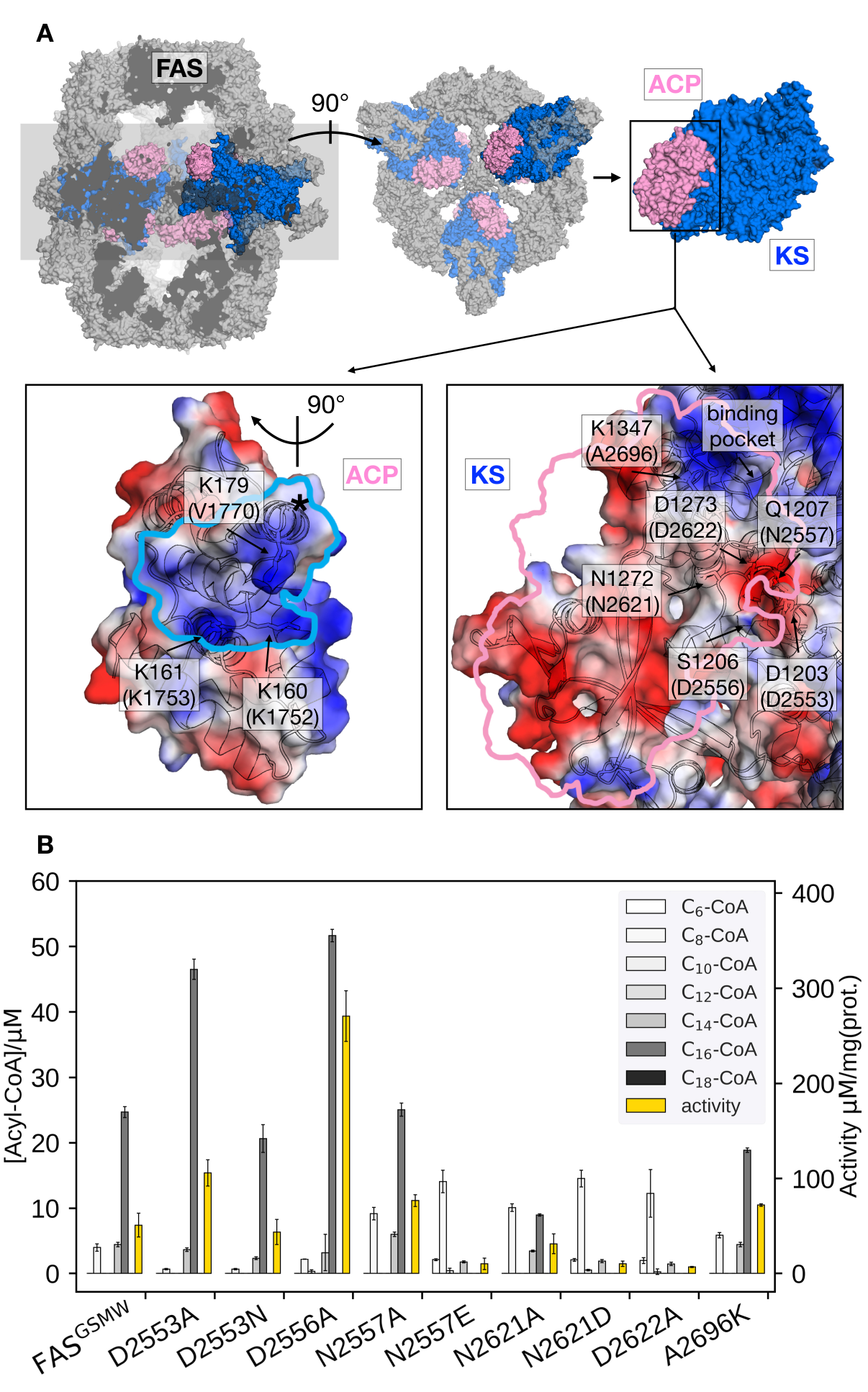
Interface engineering. (A) Structural overview of fungal FAS (pdb: 3hmj^26^) depicted as cross-section along the threefold axis and in substructures as indicated. ACP and KS are colored according to its vacuum electrostatics (generated with PyMOL; red, negatively charged; blue, positively charged). Active-site serine of ACP is highlighted (*). KS mutations are indicated in *S. cerevisiae* FAS numbering and in *C. ammoniagenes* FAS equivalent numbering in brackets. (B) Product spectra and overall enzymatic activity of FAS. KS surface mutations were introduced in FAS^GSMW^. Error bars show the standard deviation of three assays for each construct.

Our data clearly demonstrate the sensitivity of substrate shuttling in FAS to mutations in the ACP:KS interface. For several constructs, a significant increase in the fraction of short acyl-CoA esters was observed (Fig. 2B. Mutations N2557E, N2621D and D2622A were most efficient in C_8_- CoA production, yielding up to 77% C_8_-CoA of all detected CoA esters. The constructs with the highest yields in short-chain acyl-CoA were most severely impaired in their activity; i.e., N2557E, N2621D, and D2622A showed only a fraction of the specific activity of the template construct (Fig. 2C). Correlation of selective short acyl-CoA production and poor activities implies that changes in the product spectrum require the efficient suppression of specific DDIs. D2553A, D2553N and D2556A showed effects opposite to what was anticipated (Fig. 2B), and increased the fraction of long chain acyl-CoA.

We used structure-based homology modeling to gain a deeper understanding of the relation between ACP:KS domain interactions. We constructed three-dimensional models for ACP:KS using the *S. cerevisiae* FAS crystallographic structure (pdb: 2uv8) as a template (Note SI2, Fig. SI4-SI8).^17^ High crystallographic B-factors and low sequence conservation in a loop between K160 and P167 of ACP suggested local structural disorder (Fig. 3A). To account for possible flexibility in this region, we constructed two different models (A and B). In model A, we fully retained the secondary structure of the template. In model B, we locally relaxed the secondary structure restriction to account for a variation in bacterial and fungal systems (Fig. 3B). The multiple sequence alignment indicates that target loop residues V1753 and R1754 correspond to template K161 and S162, respectively (*S. cerevisiae* FAS numbering (2uv8); Fig. SI5). In *S. cerevisiae* FAS, K161 forms a salt bridge with KS D1203 that is thus not maintained in *C. ammoniagenes*, with V1753 and D2553 at the positions of K161 and D1203. R1754 in the ACP loop emerges as a possible salt-bridge partner of D2553 on KS. Model B enables this missing non-covalent interaction at the interface. After molecular dynamics (MD) relaxation, both models A and B remained close to the yeast crystal structure, with Cα backbone Root Mean Square Deviations (RMSD) of 0.74 Å and 0.93 Å, respectively (Fig. 3B). For models A and B, we performed binding-energy calculations using structures sampled in MD simulations of FAS^GSMW^ and of surface mutants, as obtained by modifying the wild type model.

**Figure 3.**
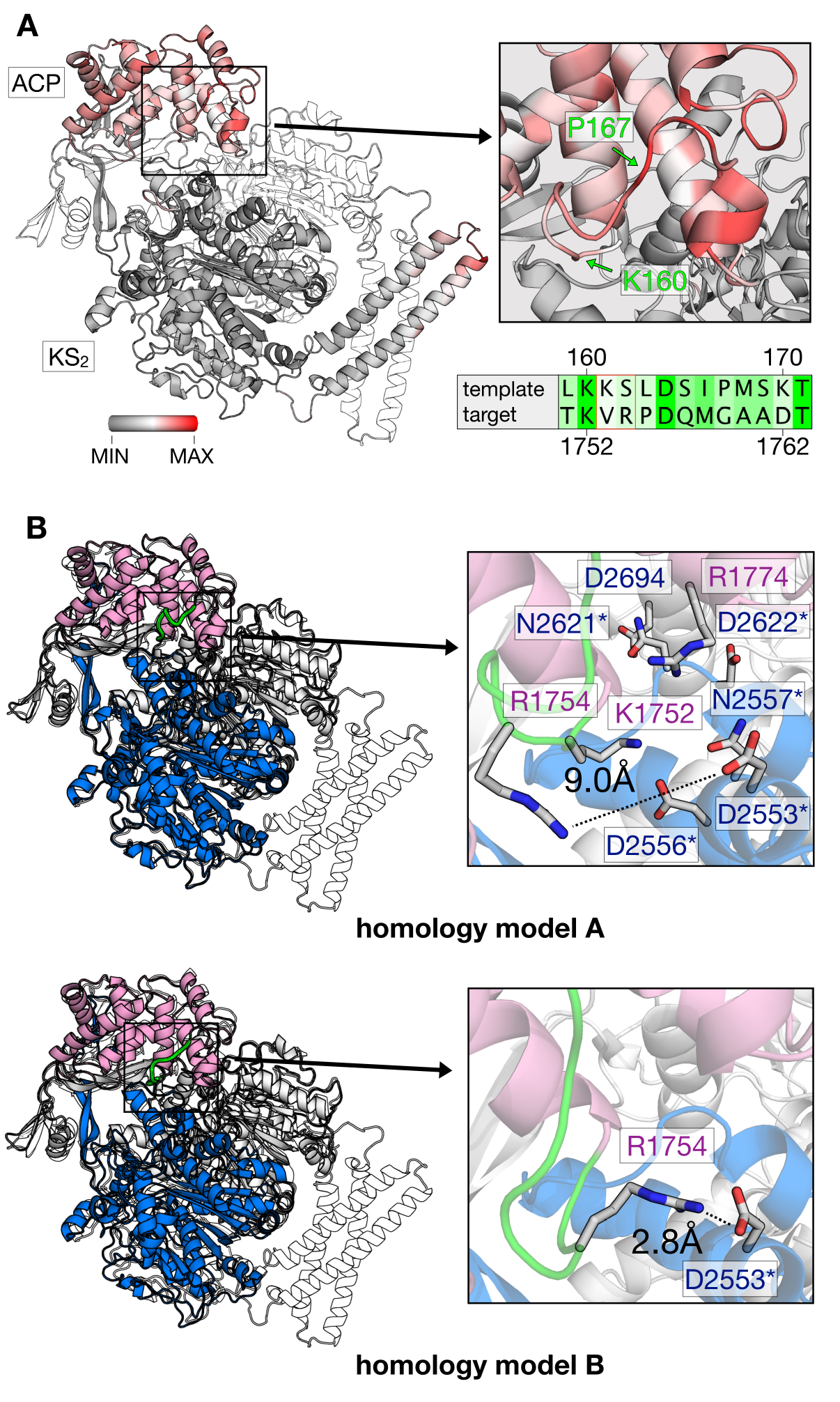
Generation of *in silico* models. (A) B factor representation of experimental template structure (B-factor range between 0 and 202). ACP binds to dimeric KS (KS_2_) as indicated (protomer colored in white). Close-up of the ACP loop between K160 and P167, and sequence alignment (green coloring reflects similarity (Fig. SI5)). (B) Structural comparison of template (*S. cerevisiae*) and target (*C. ammoniagenes*) ACP:KS complexes. *S. cerevisiae* FAS template shown in gray and *C. ammoniagenes* FAS in magenta (ACP) and blue/white (KS_2_). The ACP loop between K1752 and G1759 of ACP is highlighted in green. Models A and B differ in interactions between the interface residues R1774 and D2553.

The calculated binding affinities correlate both with the measured FAS activities and the observed product spectra (Fig. 4A). Strong ACP:KS binding is associated with the production of long acyl chains and high FAS activity. The good correlation implies that the stability of the ACP:KS interaction indeed modulates *C. ammoniagenes* FAS function, which allows us to establish detailed relations between structure, energetics, and FAS product spectrum.

**Figure 4.**
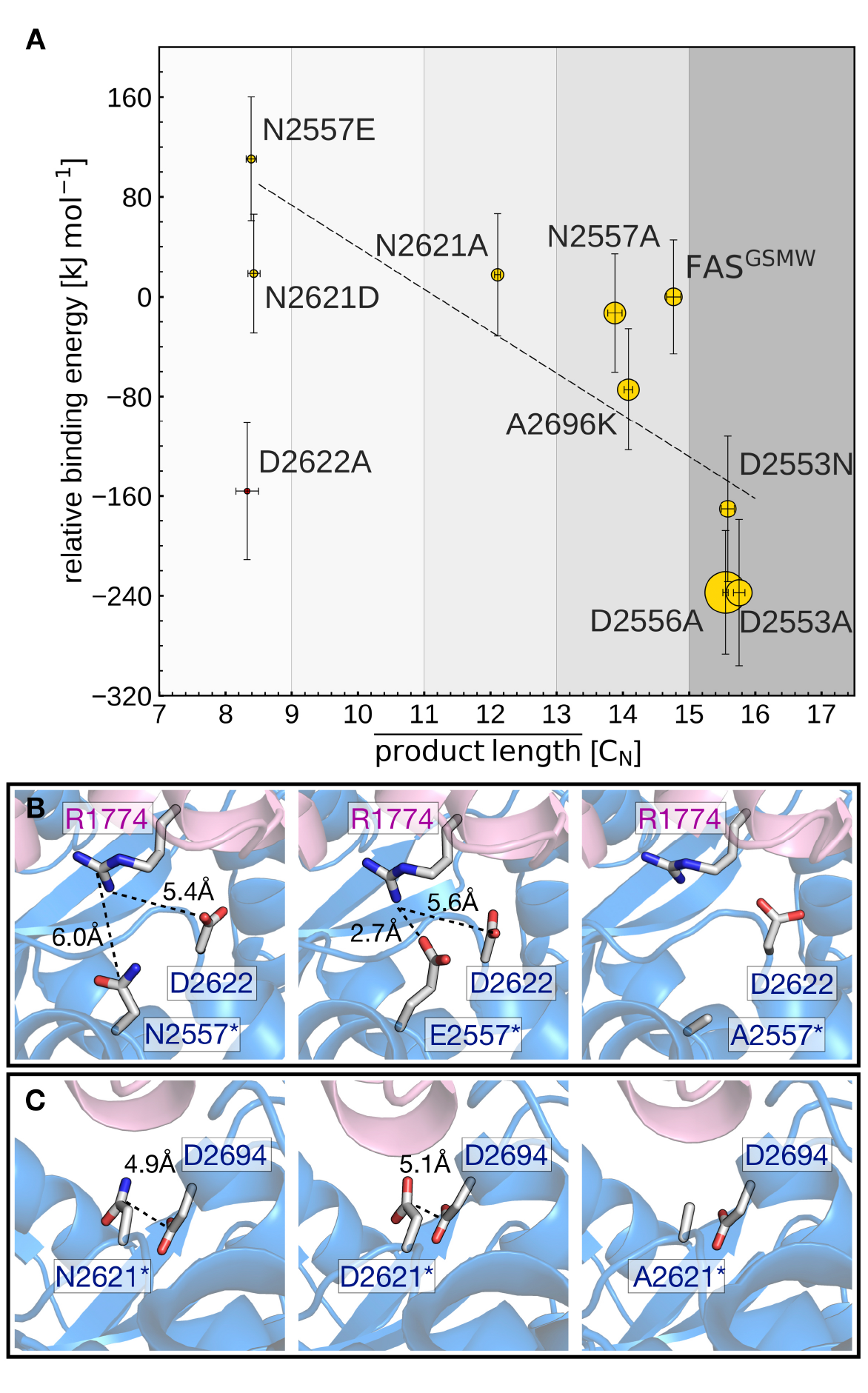
Analysis of mutated ACP:KS interfaces. (A) Calculated binding free energies (vertical axis) and measured FAS enzymatic activity (symbol radius) versus mean acyl chain length of the FAS product spectra. Error bars reflect the standard deviations from the experimental replicates (horizontal) and binding energy computations (vertical). (D) Interaction network of interface residue N2557 in the native (left) and N2557E-mutated (middle) and N2557A-mutated (right) interfaces. (E) Interface residue N2621 in the native (left) and N2621D-mutated (middle) and N2621A-mutated (right) interfaces.

Mutants N2557E and N2621D are most effective in producing C_8_-CoA and display the highest (i.e., least favorable) relative binding energies. This can be explained by mutations introducing negative charges at the DDI that affect an ionic interaction network across residues 2557, 2621, 2622, and 2694 (KS) and 1774 (ACP) (Fig. 4B and C). In the unmodified interface, the polar residue R1774 is located at distances of 6.0 Å and 5.4 Å from N2557 and D2622, respectively. By contrast, the mutant E2557 sequesters R1774 in a salt-bridge interaction. The resulting charge imbalance around D2622 weakens the polar interaction network and the interface stability (Fig. 4B). Similarly, the change in charge of N2621D disrupts the polar interaction between N2621 and D2694 (Fig. 4C). Both mutations lower FAS enzymatic activity. By contrast, mutations N2557A and N2621A do not alter the charge balance of the interface and have no effect on FAS function (Fig. 4B and C).

A2696 is located at the interface periphery and its mutations do not trigger the modulation of FAS function (Note SI3 and Fig. SI9). Mutants D2553A, D2553N and D2556A revert the FAS^GSMW^ spectrum to long chain production and display the highest (i.e., most favorable) relative binding energies. Interestingly, the energy shift of D2553A is only captured when combining the two alternative configurations of model A (the mutant) and model B (the wild-type). In such a scheme, the polarity of position 2553 (KS) influences the orientation of R1754 (ACP) that forms a salt-bridge with D2553 and is repelled by A2553 (Note SI4 and Fig. SI10). Mutant D2553N, despite a possible change in charge, has a minor impact on R1754 orientation (Note SI4 and Fig. SI10). In D2556A, the negative charge deletion leads to a minor but effective structural reorganization of the polar network at the interface (Note SI5 and Fig. SI11). In case of the charged-to-neutral mutation D2622A at the center of a network of ionic interactions, the limited sampling of motions in the computation of binding affinities is likely insufficient to capture a major reorganization of the interface, providing an explanation for this single outlier (Note SI6 and Fig. SI12).

As a further challenge to modeling substrate shuttling, binding energies are not the only determinant of DDI kinetics. Effects of accessibility and confinement will impact the frequency and duration of productive interaction between ACP and KS. As a first step towards capturing substrate shuttling kinetics, we set up coarse-grained simulations to count frequencies of competitive binding events of ACP to wildtype and surface mutated KS (Note SI7 and Fig. SI13).^19^ The kinetic approach did, however, not lead to conclusive data, reflecting both the challenges in simulating binding kinetics in a confined environment and the paucity of experimental information required for model generation.

As its main result, this study shows that single mutations at domain-domain interfaces can severely perturb substrate shuttling in FASs. *In silico* modeling demonstrates that mutations are particularly invasive when inducing charge imbalances at the interfaces, which agrees with earlier finding on the importance of electrostatic complementarity of ACP:catalytic domains interfaces.^5,17,20^ Engineering of DDIs proved to be powerful in modulating the chain length spectrum of *C. ammoniagenes* FAS, and some mutations, i.e., N2557E, N2621D and N2622A, produced C_8_-CoA with high selectivity. However, the strong decrease in activity of the mutated FASs questions a true biotechnological relevance of DDI engineering in FASs, and rather illustrates an inherent “low-resolution” problem of the approach. While engineering of substrate binding channels^12^ or the thioesterase-mediated hydrolyzation of acyl-ACP of certain chain length^13,21–22^ hijacks FA synthesis at a specific acyl-ACP chain length, ACP surface mutations interfere already in initial FA cycles, causing the overall drop in activity.

The observed sensitivity of substrate shuttling is also interesting for the related PKSs. Similar to FASs, engineering of substrate shuttling may enable modulation of product synthesis in iterative polyketide synthases (PKSs).^23^ In modular PKSs, ACPs are involved in the substrate shuttling within one multienzyme complex (module), but also in the translocation of the substrate to a downstream module. The strong influence by even single mutations implies that a successful assembly of such modules for the design of new chimeric biosynthetic pathways will strongly rely on the effective adaptation of interfaces across the borders of the chimeric assembly lines.^24^

## ASSOCIATED CONTENT

### Supporting Information

Supporting Information is available.

### Notes

Concerning competing financial interest, J.G. and M.G. declare that they are inventors of two patents concerning the production of short fatty acids (EP patent application 15 162 192.7 filed on April 1st, 2015 and EP patent application 15 174 342.4 filed on June 29th, 2015).

## ACKNOWLEDGMENT

This work was supported by a Lichtenberg grant of the Volkswagen Foundation to M.G. (grant number 85701). Further support was received by the LOEWE program (Landes-Offensive zur Entwicklung wissenschaftlich-ökonomischer Exzellenz) of the state of Hesse conducted within the framework of the MegaSyn Research Cluster and by the Max Planck Society. E.R. acknowledges Dr. Ramachandra Bhaskara for many useful inputs and valuable comments.

